# Molecular Crowding Facilitates Bundling of IMPDH Polymers and Cytoophidium Formation

**DOI:** 10.1101/2022.03.15.484061

**Authors:** Chia-Chun Chang, Min Peng, Jiale Zhong, Ziheng Zhang, Gerson Dierley Keppeke, Li-Ying Sung, Ji-Long Liu

## Abstract

The cytoophidium is a unique type of membraneless compartment comprising of filamentous protein polymers. Inosine monophosphate dehydrogenase (IMPDH) catalyzes the rate-limiting step of *de novo* GTP biosynthesis and plays critical roles in active cell metabolism. However, the molecular regulation of cytoophidium formation is poorly understood. Here we show that human IMPDH2 polymers bundle up to form cytoophidium-like aggregates in vitro when macromolecular crowders are present. The self-association of IMPDH polymers is suggested to rely on electrostatic interactions. In cells, the increase of molecular crowding with hyperosmotic medium induces cytoophidia, while the decrease of that by the inhibition of RNA synthesis perturbs cytoophidium assembly. In addition to IMPDH, CTPS and PRPS cytoophidium could be also induced by hyperosmolality, suggesting a universal phenomenon of cytoophidium-forming proteins. Finally, our results indicate that the cytoophidium can prolong the half-life of IMPDH, which is proposed to be one of conserved functions of this subcellular compartment.

## Introduction

Membraneless organelles, such as processing body (P-body), Cajal body and stress granules, are special compartment of proteins and/or RNAs, and responsible for diverse biological functions of the cell. Among them, a unique type of protein aggregates, the cytoophidium (cellular snake in Greek), is formed by large bundles of filamentous polymers of functional proteins (Liu, 2016). The cytoophidium initially designates a distinctive compartment of CTP synthase (CTPS) in *Drosophila* tissues (Liu, 2010). Soon later, some studies have discovered similar filament-forming properties of other metabolic enzymes and applied the name to novel intracellular filaments in various organisms, such as asparagine synthase (ASNS) cytoophidium, pyrroline-5-carboxylate synthase (P5CS) cytoophidium and inosine monophosphate dehydrogenase (IMPDH) cytoophidium (Carcamo et al., 2011; Chang et al., 2015; Zhang et al., 2020; Zhang et al., 2018).

In mammalian models, CTPS and IMPDH are the best-known cytoophidium-forming enzymes. Previously, the polymer structures of CTPS and IMPDH have been resolved by cryo-EM (Anthony et al., 2017; Johnson and Kollman, 2020; Lynch et al., 2017; Lynch and Kollman, 2020). The CTPS polymer is composed by CTPS tetramers and able to moderate the end-product inhibition by CTP binding (Lynch et al., 2017; Zhou et al., 2019). Similarly, IMPDH octamers can stack back-to-back to form filaments and thereby enhance the activity in the presence of the allosteric inhibitor GTP (Anthony et al., 2017; Johnson and Kollman, 2020). Forming the cytoophidium can also attenuate CTPS ubiquitination and protect CTPS from proteasomal degradation (Lin et al., 2018; Sun and Liu, 2019a). Assembly of both CTPS and IMPDH cytoophidium has been shown positively correlated with mTORC activity, active glycolysis and rapid cell proliferation in cell types including lymphocytes and certain cancers, suggesting their physiological importance (Calise et al., 2018; Chang et al., 2015; Duong-Ly et al., 2018; Keppeke et al., 2019; Keppeke et al., 2018; Peng et al., 2021; Sun and Liu, 2019b).

IMPDH catalyzes the conversion of IMP to XMP, which is the rate-limiting step of *de novo* GTP biosynthesis. The precise regulation of IMPDH activity is critical in the coordination of metabolic pathways and cellular status. Specific point mutations on IMPDH1 and IMPDH2 can result in retinopathy and neuropathy, respectively (Bowne et al., 2006; Burrell and Kollman, 2022; Zech et al., 2020). More recently, researchers have linked the elevation of IMPDH expression and activity with the increase of rRNA and tRNA synthesis, which promotes tumor progression (Kofuji et al., 2019). Meanwhile, an increased number of cells with IMPDH cytoophidia has been observed in acral melanomas (Keppeke et al., 2019). Therefore, IMPDH has long been considered as a promising drug target for autoimmune diseases, viral infection and some cancers (Hedstrom, 2009).

Given accumulative evidences implying the significance of IMPDH cytoophidium in areas including rheumatology, immunology and oncology, we seek to investigate the underlying mechanisms of IMPDH cytoophidium formation.

In this study, we show that IMPDH polymers can self-associate into large bundles in the presence of molecular crowders. The negative charge at the loop^214-217^ of human IMPDH2 may play a critical role in interpolymer interactions. In cells, IMPDH cytoophidium assembly is initiated with the formation of an amorphous clump, which is in association with the ER and can rapidly transform into filaments. The aggregation of IMPDH polymers is regulated by the molecular crowding. While the reduction of intracellular macromolecules with the inhibition of RNA synthesis attenuates drug-induced IMPDH cytoophidia, the increase of molecular crowding with hyperosmotic medium stimulates the formation of CTPS, PRPS and IMPDH cytoophidium. In the cells with cytoophidia, the IMPDH display longer half-life, suggesting that the cytoophidium protects its component proteins from degradation. Our findings reveal molecular mechanisms of the regulation and formation of IMPDH cytoophidium and imply the general function of this cellular compartment.

## Results

### Molecular crowders trigger in vitro bundling of IMPDH polymers

The cytoophidium in micron-scale size is demonstrated to be a large bundle of filamentous polymers (Chakraborty et al., 2020; Ingerson-Mahar et al., 2010; Thomas et al., 2012). We have previously shown that CTPS and IMPDH form separate cytoophidia in the cell (Chang et al., 2018; Chang et al., 2015). Although interactions between two cytoophidia may present, CTPS and IMPDH polymers do not mix up within the same micron-scale filament. These suggest that the interactions holding filamentous polymers together have certain specificity.

Filamentous polymers of the autosomal dominant retinopathy 10 (adRP10) related mutant human IMPDH1^D226N^ recombinant can spontaneously aggregate into bundles in vitro, while the wildtype protein can only form separate polymers under the same condition (Labesse et al., 2013). However, the presence of macromolecular crowder Ficoll-70 triggers further aggregation of wildtype human IMPDH2 polymers, implying that cytoophidium assembly may solely rely on the interaction between polymers, which could be enhanced by the increase of molecular crowding (Fernandez-Justel et al., 2019).

To test this notion, we attempted to reconstitute IMPDH cytoophidium in vitro with human IMPDH2 recombinant protein and the supplementation of molecular crowder polyethylene glycol (PEG-4000). When pure human IMPDH2 protein was examined with negative staining, many polymers were observed (Figure 1A). Additional ATP (100 μM) in the mixture considerably increased the amount and the length of polymers, but all polymers remained separate (Figure 1B). Strikingly, when the PEG-4000 was added into the mixture at the concentration of 100 mg/ml, microns long bundles were observed (Figure 1C). The increase of PEG concentration did not enlarge the bundles, but rendered large networks of tangled bundles (Figure 1D).

**Figure 1.**
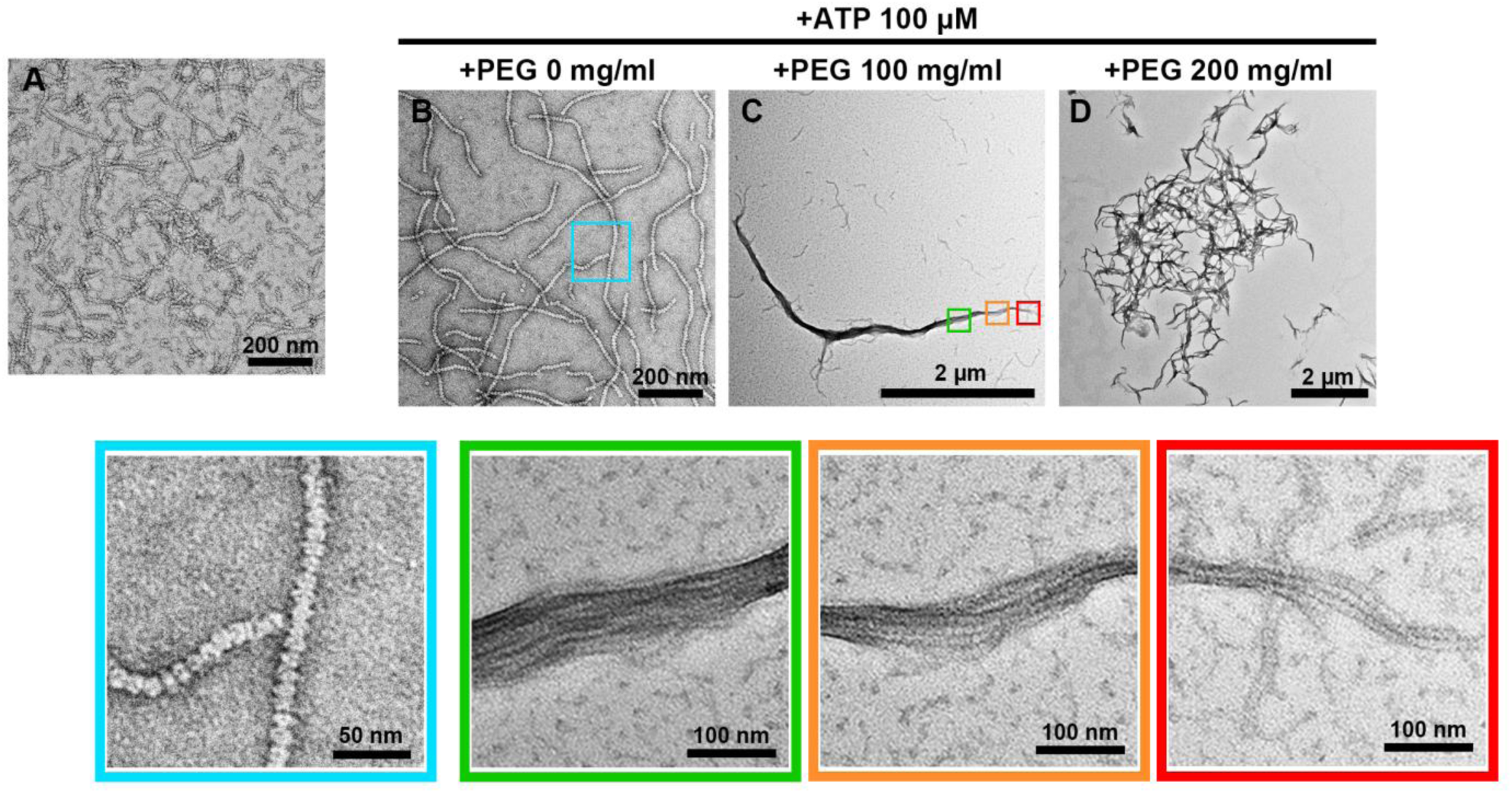
Molecular crowder PEG-4000 promotes in vitro reconstitution of IMPDH cytoophidium. (A-D) Negative staining images of hIMPDH2 recombinant protein incubated with different conditions for 1 hour. (A) hIMPDH2 protein was incubated in the buffer without the supplementation of ligands. (B-D) hIMPDH2 protein was incubated in the buffer containing 100 μM ATP and PEG-4000 at 0, 100, 200 mg/ml. Lower panels show magnified images of corresponding areas in (B) and (C).

### IMPDH polymers interact through electrostatic interactions

The formation of filamentous aggregates has been proposed to be driven by electrostatic interactions and the steric compatibility of such protein filaments (Petrovska et al., 2014). Electrostatic interactions are relatively weak and transient interactions that are regulated by salt, pH, post-translational modifications and the protein concentrations (Dumetz et al., 2008).

We analyzed the electrostatic surface potential of human IMPDH2 polymer with a structural model (PDB ID: 6U8N) and found the loop^214-217^, which comprises of two aspartate and two glutamate, at the Bateman domain displays intense negative charge (Figure 2A-C). The sequence comparison between different IMPDH isoforms of human, mouse and zebrafish, in which species the filamentation of IMPDH has been reported, shows that at least two of these four residues are with negative charge in all IMPDH sequences (Figure 2D). To assess whether the negative charge at the loop^214-217^ is required for the inter-polymer interactions within the cytoophidium, we replaced all four residues with alanine (hIMPDH2^4A^). Subsequently, we induced cytoophidia in HEK 293T cells overexpressing wildtype hIMPDH2 or hIMPDH2^4A^ with mycophenolic acid (MPA) for 2 hours. MPA is an IMPDH specific inhibitor and can effectively stimulate IMPDH cytoophidium formation in many cell types and organisms (Carcamo et al., 2011; Ji et al., 2006; Keppeke et al., 2015; Keppeke et al., 2021). As the result, while long IMPDH filaments were observed in most cells overexpressing wildtype hIMPDH2, most hIMPDH2^4A^ expressing cells could only form small clumps (Figure 2E and F), suggesting that negative charge at the loop^214-217^ participates in the interaction within the filament. Notably, since the endogenous wildtype IMPDH proteins were also present in transfected cells, patterns of cytoophidia could be affected by the ratio of wildtype and mutant IMPDH.

**Figure 2.**
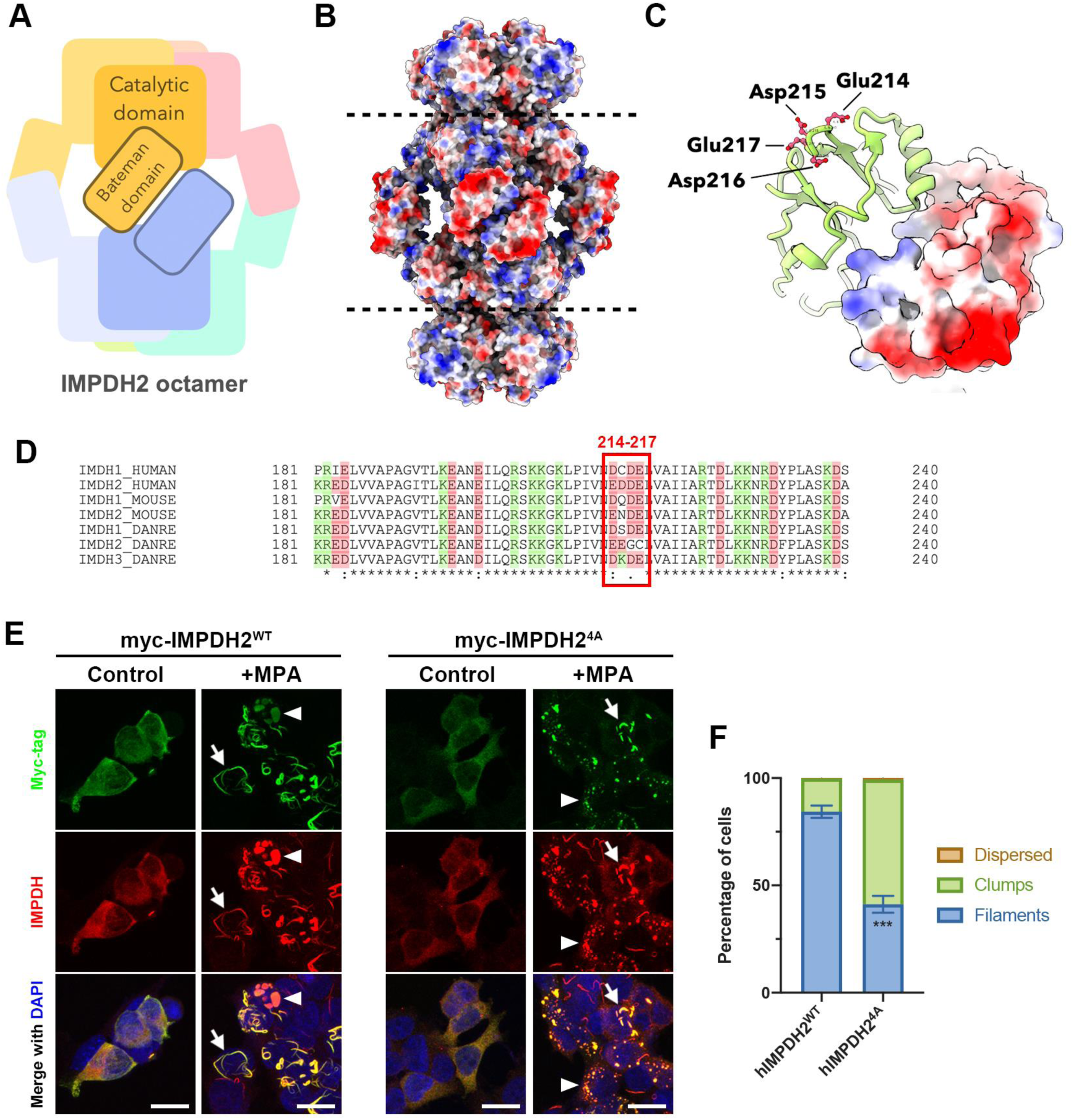
The mutation at loop^214-217^ of human IMPDH2 disturbs the cytoophidium assembly in cells. (A) Illustration of the IMPDH octamer. (B) Electrostatic surface model of hIMPDH2 polymer. (C) Electrostatic surface model of the Bateman domain of hIMPDH2. Residues at loop^214-217^ are indicated. (D) The sequence comparison of the CBS subdomain of human, mouse and zebrafish IMPDH isoforms. Residues with positive and negative charge are highlighted in green and red, respectively. (E) Immunofluorescence of HEK 293T cells overexpressing myc-hIMPDH2^WT^ and myc-hIMPDH2^4A^. Cells were treated with MPA for 2 hours before fixation. Cells displaying IMPDH filaments are indicated with arrows and cells displaying IMPDH clumps are indicated with arrowheads. Scale bars = 20 μm. (F) Quantification of cells displaying different IMPDH patterns in (E). Error bars = S.E.M. Student’s *t* test, \*\*\**p* < 0.001.

### Cytoophidium assembly goes through a state transition

In order to understand the initiation of cytoophidium assembly, we captured the early phases of the process. In wildtype HeLa cells, small IMPDH cytoophidia could be found in cells in just a few minutes upon the MPA induction (Figure 3A). Surprisingly, some IMPDH aggregates without clear filamentous appearance were observed in a small portion of cells treated with MPA for less than 5 minutes (Figure 3A). In an OFP-IMPDH2 overexpressing HeLa cell line, many OFP-IMPDH clumps were intertwined with or surrounded by the ER (Figure 3B-E). Intriguingly, these OFP-IMPDH2 clumps can remain in the cell for more than an hour in some cases (Figure 3B). We then performed live-cell imaging on OFP-IMPDH2 overexpressing cells to observe the dynamics of these IMPDH clumps. These amorphous IMPDH clumps can migrate, fuse with one another and split in association of ER dynamics (Figure 3E and F). Occasionally, we observed the transition of a clump from its amorphous state into a filament, showing that the such an amorphous IMPDH clump is the precursor of the filamentous cytoophidium (Figure 3G). Since the electrostatic interaction requires precise pairing of specific domains of the proteins, we suspect that the bulky fluorescent tag may destabilize inter-polymer interactions and thereby delay their condensation and transformation. The same phenomenon has not been not observed when the fluorescent tag was replaced by small tags.

**Figure 3.**
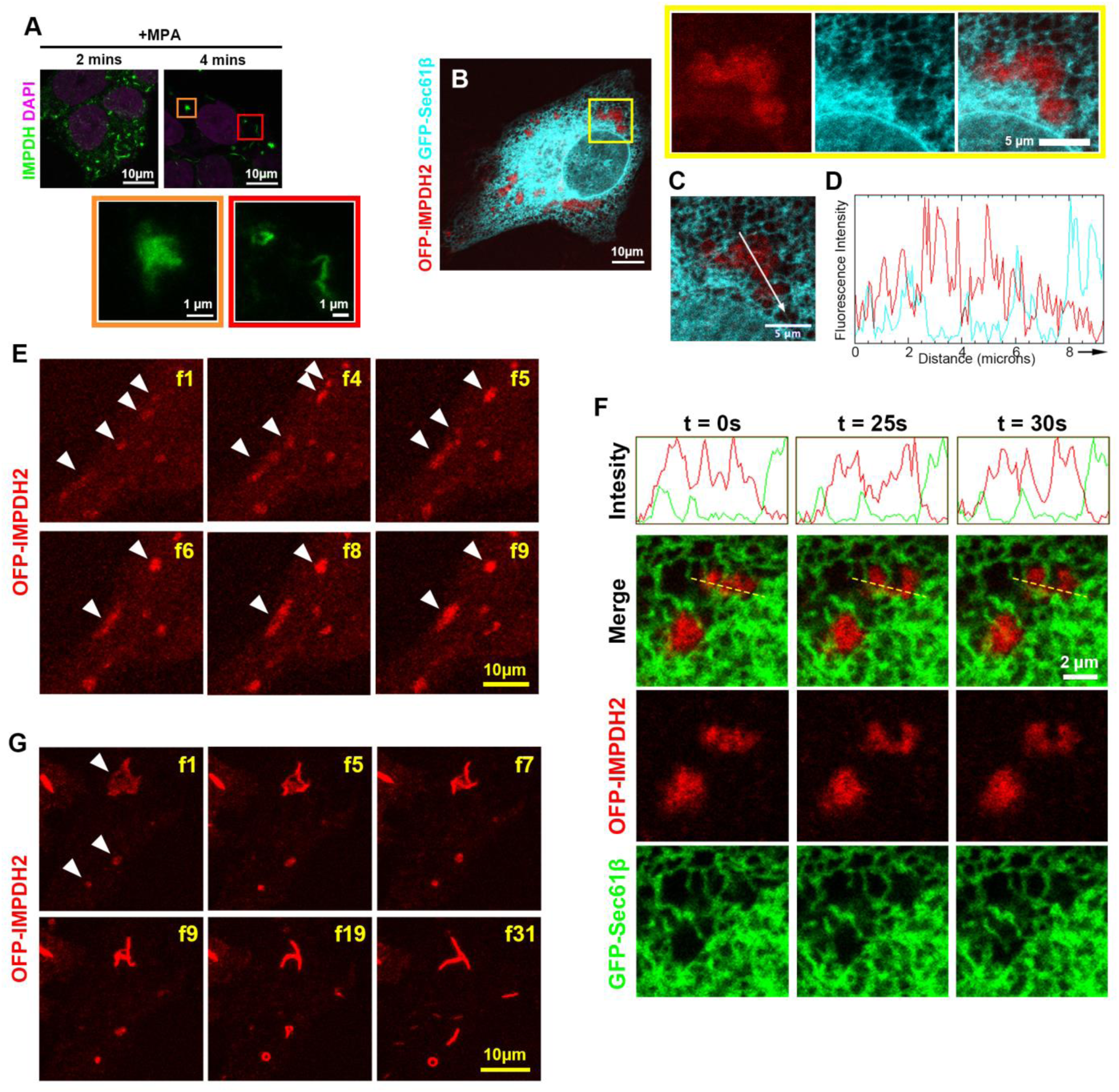
Amorphous IMPDH clump is the precursor state of the filamentous cytoophidium. (A) Immunofluorescence of wildtype HeLa cells treated with MPA for 2 and 4 minutes, respectively. Magnified images of an amorphous IMPDH clump (orange box) and filamentous cytoophidia (red box) are corresponding to selected areas in (A). (B) Images of HeLa cells expressing OFP-IMPDH2 and GFP-Sec61β fusion proteins. Cells were treated with MPA for 1 hour. Magnified images show an amorphous IMPDH clump surrounded by the ER in selected areas. (C) Single plane image of the IMPDH clump shown in magnified images in (B). (D) Fluorescent intensity of OFP-IMPDH2 and GFP-Sec61β signals in (C). The *x* axis corresponds to the direction and area of measurement indicated by the arrow in (C). (E-G) Representative frames of live-cell imaging of OFP-IMPDH2 expressing HeLa cells treated with MPA. (E) Selected frames of movie S1 showing the movement and fusion of IMPDH clumps (arrowheads). (F) Selected frames of movie S2 showing an IMPDH clump split in association with ER tubule dynamics. The dashed line corresponds to the area of fluorescence intensity measurement. (G) Selected frames of movie S3 showing the clump-to-filament transition. Time intervals of each frame is 40 sec in (E) and 60 sec in (G).

Therefore, we hypothesize that the assembly of IMPDH cytoophidium is initiated with the loose connections between filamentous polymers, which results in an amorphous state. Subsequent transformation into the filaments would be achieved by the accumulation of interactions between long polymers. Such interactions may be regulated by concentrations of IMPDH polymers and local macromolecules. The dynamics of membrane-bound organelles, such as ER, may also contribute to the regulation.

### Hyperosmolality triggers rapid and reversible cytoophidium assembly

To test if the formation of the IMPDH cytoophidium is controlled by molecular crowding in the cell, we treated the cells with hyperosmotic medium, which would rapidly dehydrate the cells, thereby increasing concentrations of intracellular solutes. HEK 293T cells were cultured in the medium containing sucrose at concentrations ranging from 25 mM to 300 mM for one hour before fixation. While no enrichment of cytoophidia was observed in cells treated with 25 mM and 50 mM sucrose, medium with 100 mM and 150 mM sucrose induced mature cytoophidia in 18.9% and 54.7% of cells, respectively (Figure 4A and B). Other cell lines, including HeLa cells, MCF7 cells and HCT116 cells also exhibited increasing numbers of IMPDH cytoophidia in a dose-dependent manner, showing that cytoophidium assembly induced by hyperosmotic medium is a general phenomenon in human cells (Figure 4C and D).

**Figure 4.**
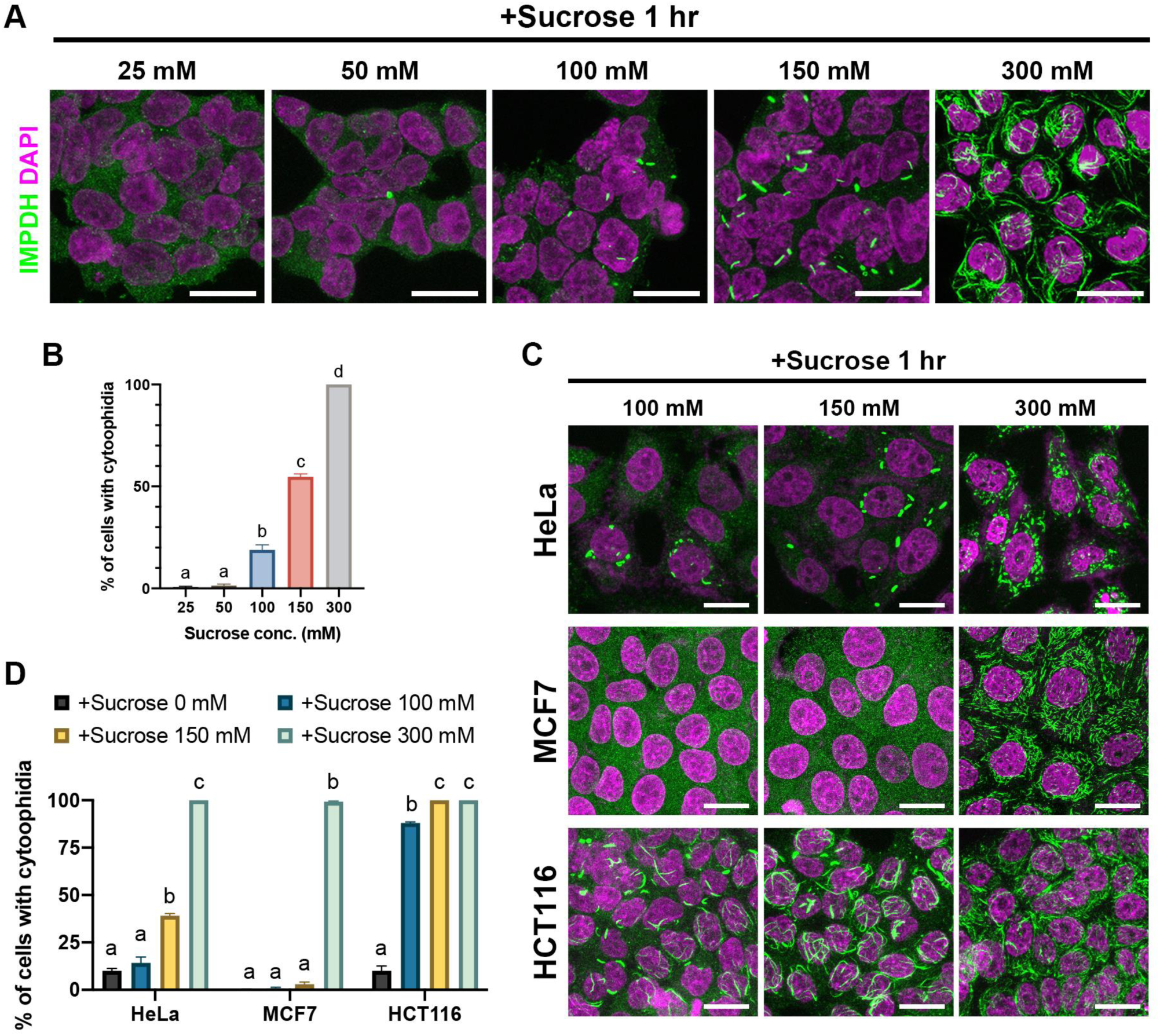
Hyperosmotic medium induces IMPDH cytoophidium assembly in multiple cell lines. (A) Immunofluorescence of wildtype HEK 293T cells treated with sucrose at different concentrations for 1 hour. (B) Quantification of the percentage of cells with cytoophidia under the treatments shown in (A). (C) Immunofluorescence of wildtype HeLa, MCF7 and HCT116 cells treated with sucrose at different concentrations for 1 hour. (D) Quantification of the percentage of cells with cytoophidia under the treatments shown in (C). Scale bars = 20 μm in all panels. Error bars = S.E.M. Tukey’s test was used in the comparison in (B) and (D).

Interestingly, nearly all cells treated with hyperosmotic medium with 300 mM sucrose displayed IMPDH cytoophidia (Figure 4). It is known that the formation of protein dimer, oligomer or polymer is preferable in the crowded milieu in order to minimize the excluded volume and overall crowding (Ralston, 1990). Rapid and reversible formation of large protein aggregates and membraneless organelles could be induced by hyperosmolality (Jalihal et al., 2020). When HEK 293T cells were treated with 300 mM sucrose, a great amount of small IMPDH filaments appeared in all cells within 3 minutes of culture (Figure 5A). These IMPDH cytoophidia will undergo elongation and fusion, eventually forming a few large ones (Figure 5A). Conversely, when the medium was replaced with isosmotic medium, these large cytoophidia completely disassociated within 5 minutes, indicating that hyperosmolality-induced IMPDH cytoophidium formation is also a rapid and reversible process (Figure 5B).

**Figure 5.**
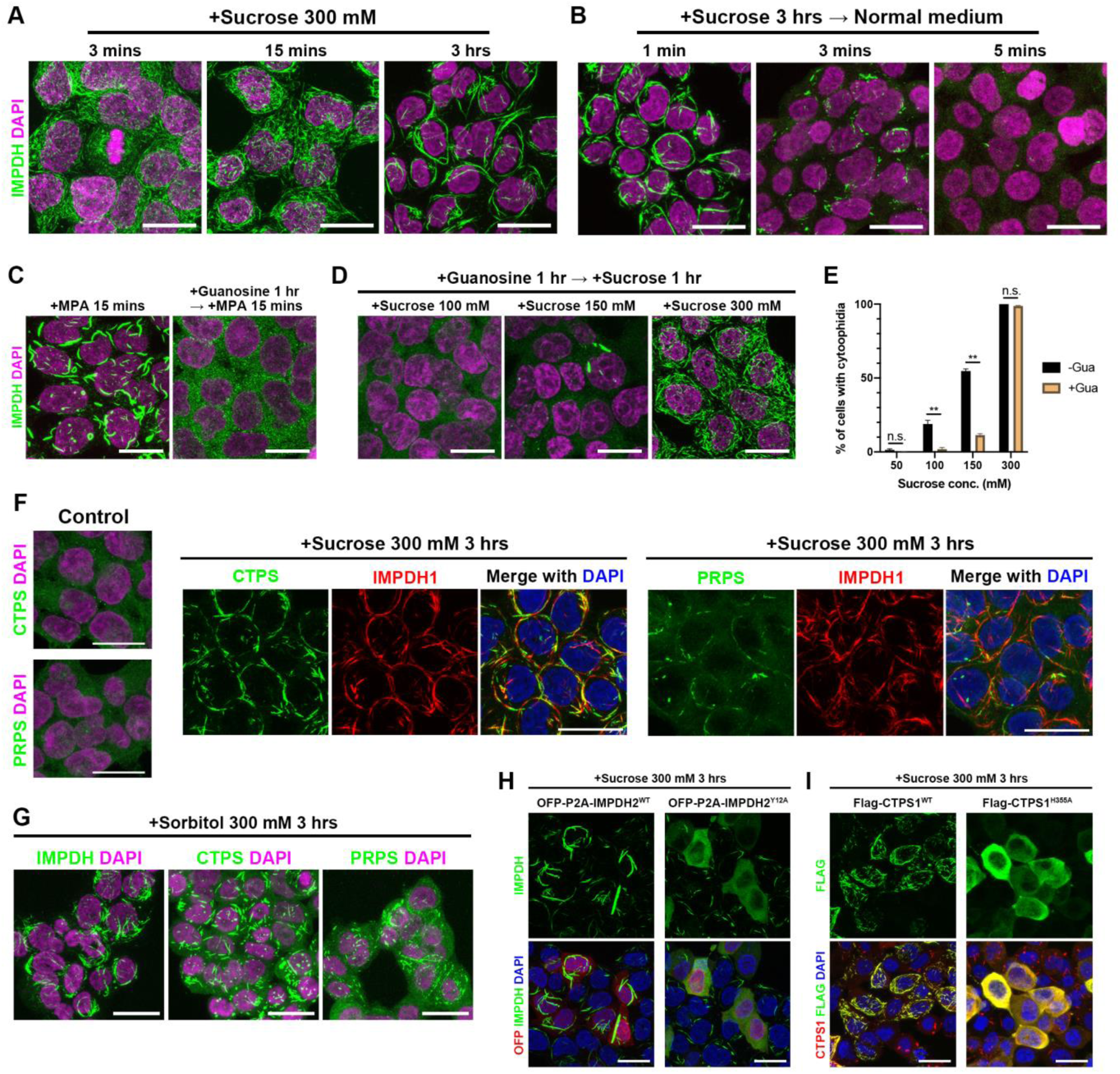
Hyperosmotic medium induces the protein polymerization and aggregation of filamentous polymers. (A) Immunofluorescence of wildtype HEK 293T cells treated with 300 mM sucrose for different periods of time. (B) Immunofluorescence of wildtype HEK 293T cells pre-treated with 300 mM sucrose for 3 hours then treated with isosmotic medium for 1 to 5 minutes. (C) and (D) Immunofluorescence of wildtype HEK 293T cells pre-treated with 100 μM guanosine for 1 hour and then treated with 100 μM MPA (C) or sucrose (D). (E) Quantitative data of the presence of hyperosmolality-induced cytoophidia in HEK 293T cells pre-treated with and without guanosine. (F) Immunofluorescence of wildtype HEK 293T cells for hyperosmolality-induced CTPS, PRPS and IMPDH cytoophidium. (G) Immunofluorescence of wildtype HEK 293T cells for CTPS, PRPS and IMPDH cytoophidium in the medium containing 300 mM sorbitol. (H) and (I) Immunofluorescence for IMPDH and CTPS in HEK 293T cells expressing OFP-P2A-IMPDH2^WT^, OFP-P2A-IMPDH^Y12A^, Flag-CTPS1^WT^ and Flag-CTPS1^H355A^. Scale bars = 20 μm in all panels. Error bars = S.E.M. Student’s *t* test, \*\**p* < 0.01 in (E).

The polymerization of IMPDH octamers is controlled by the conformational changes, which is determined by the binding of ATP and GTP (Anthony et al., 2017; Johnson and Kollman, 2020). Consistently, IMPDH cytoophidia in cells could be disrupted by elevating intracellular GTP levels with the treatment of guanosine or GTP (Ji et al., 2006; Keppeke et al., 2018). When HEK 293T cells were pre-treated with guanosine, the MPA-induced cytoophidium assembly was prohibited (Figure 5C). However, the same pre-treatment failed to prevent IMPDH filamentation induced by hyperosmotic medium with 300 mM sucrose, suggesting that filament-forming proteins may undergo polymerization regardless the ordinary regulation when cytoplasmic crowding reaches certain levels (Figure 5D and E).

Polymerization and cytoophidium-forming properties have been revealed in an increasing number of enzymes over the last decade. Since the excluded volume effect is a general physical principle of macromolecules, IMPDH cytoophidium is unlikely the only protein filament induced by hyperosmolality. In mammalian models, CTPS and PRPS are two other cytoophidium-forming enzymes supported by multiple studies (Begovich et al., 2020; Gou et al., 2014; Lin et al., 2018; Noree et al., 2019). By performing the immunostaining on HEK 293T cells treated with hyperosmotic medium for 3 hours, we found CTPS and PRPS cytoophidia in most of cells (Figure 5F). The treatment of 300 mM sorbitol was applied to the cells as the sucrose substitute and that also induced these three types of cytoophidia in most cells (Figure 5G). The localization of some other enzymes, including PAICS, GMPS and HPRT in these cells were also examined but no filamentous structure was found (Figure S1).

The polymer structures of IMPDH2 and CTPS1 have been resolved and Y12A of IMPDH2 and H355A of CTPS1 mutations were employed to disrupt the interaction between neighboring protomers in previous studies (Anthony et al., 2017; Johnson and Kollman, 2020; Lynch et al., 2017). To test if the formation of filamentous polymers is still a prerequisite of assembling large filaments under hyperosmolality, HEK 293T cells were transfected with constructs encoding OFP-P2A-IMPDH2^Y12A^ or Flag-CTPS1^H355A^ mutant proteins and their wildtype counterparts before being treated with hyperosmotic medium. While cells overexpressing wildtype IMPDH2 showed large cytoophidia under the treatment, dispersed patterns of IMPDH were observed in most cells expressing mutant IMPDH2 (Figure 5H). Similarly, CTPS cytoophidium assembly under this condition was impaired in cells overexpressing mutant CTPS1 (Figure 5I). These suggest that only filamentous polymers can assemble filamentous aggregates under hyperosmotic conditions.

We also treated GFP-hIMPDH2^WT^ overexpressing HEK 293T cells with hyperosmotic medium in order to observe the formation of cytoophidia in live cells. However, only aggregates in the appearance of dots or short speckles were observed, suggesting that the inter-polymer interactions were interfered by the bulky tags in hyperosmotic conditions (Figure S2)

### Inhibition of RNA synthesis perturbs IMPDH cytoophidium assembly

Ribosomes are one of the most abundant macromolecules in the cell and serve as crowders for tuning the cytoplasmic viscosity and the effective diffusion coefficient of macromolecules (Delarue et al., 2018). We treated HEK 293T cells with CX-5461, an inhibitor of RNA polymerase I, for 3 hours prior to the supplementation of MPA for 15 and 30 minutes. The rRNA synthesis was monitored with the incorporation of 5-ethynyl-uridine (EU) in nucleoli, which labels newly synthesized RNAs (Figure 6A and C). The amounts of cytoophidia in size exceeding a threshold (> 3 pixel^2^) were quantified. Significantly fewer cytoophidia were found in cells upon 15 minutes of MPA induction (Figure 6A). However, no significant difference was shown when the duration of MPA treatment was extended to 30 minutes (Figure 6A and C), suggesting that the inhibition of rRNA synthesis delays but not prohibits IMPDH cytoophidium assembly.

**Figure 6.**
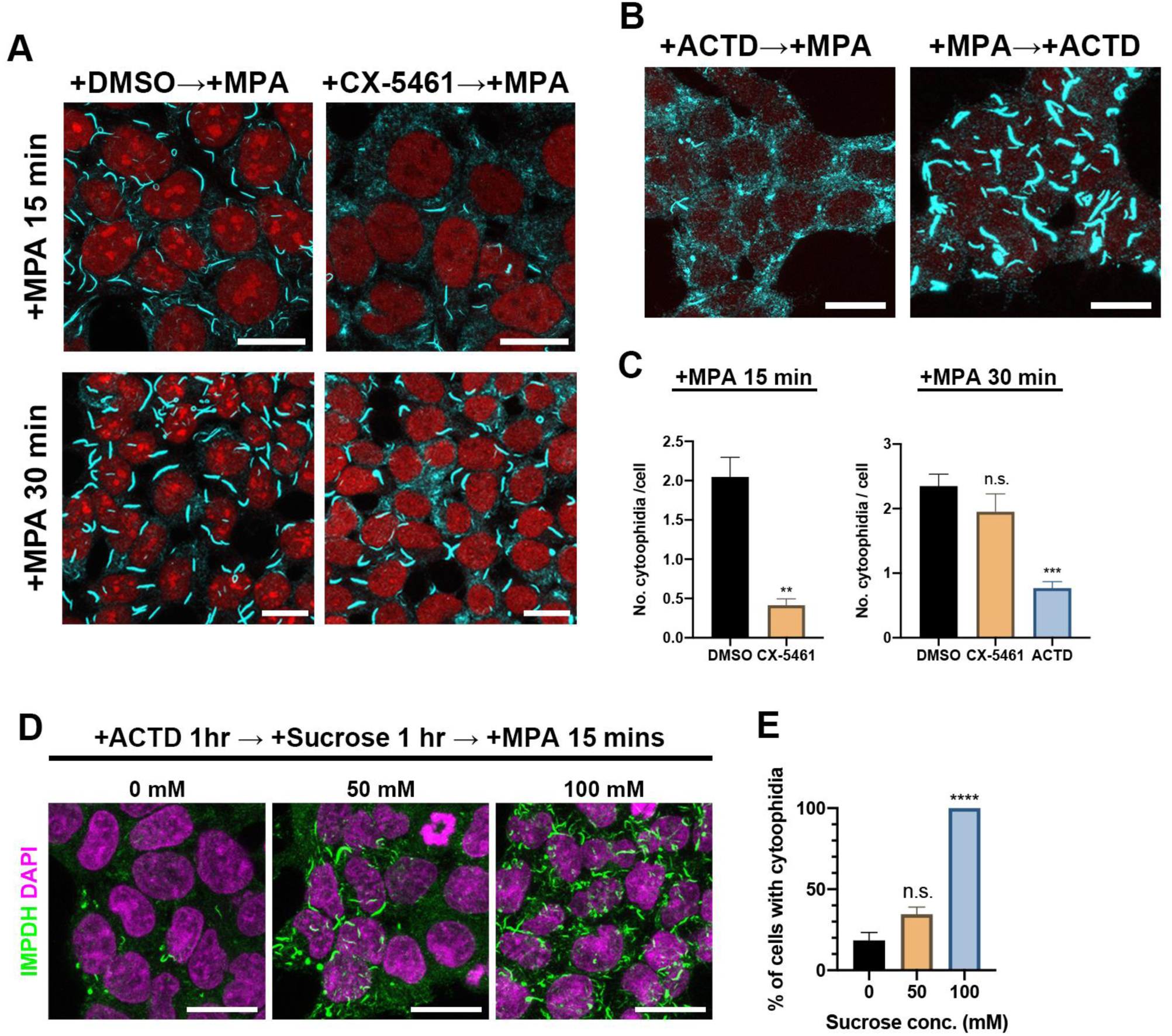
Inhibition of transcription perturbs the MPA-induced IMPDH cytoophidium assembly. (A) Immunofluorescence of wildtype HEK 293T cells treated with DMSO or 1 μM CX-5461 for 3 hours prior to 15 and 30 minutes of MPA treatment. (B) Immunofluorescence of wildtype HEK 293T cells treated with 1 μM ACTD for 1hour before or after the treatment of MPA. (C) Quantitative data of the abundance of IMPDH cytoophidia under different treatments shown in (A) and (B). (D) Immunofluorescence of wildtype HEK 293T cells treated with ACTD for 1 hour and sucrose for another 1 hour prior to cytoophidium induction with MPA. (E) Quantitative data of the proportion of cells with cytoophidia in the conditions shown in (D). Scale bars = 20 μm in all panels. Error bars = S.E.M. Student’s *t* test, \*\**p* < 0.01, *** *p* < 0.001, **** *p* < 0.0001 in (C) and (E).

Next, we wondered if the inhibition of global transcription, which should reduce the concentrations of total RNAs, ribosomes and proteins in the cell, would render more prominent defects on cytoophidium formation. HEK 293T cells were treated with actinomycin D (ACTD), a compound prevents the elongation of RNA chains, for 1 hour before the supplementation of MPA. After 30 minutes of MPA induction, significantly fewer cytoophidia were observed (Figure 6B and C). However, such a reduction of cytoophidia was not seen in cells pre-treated with MPA prior to ACTD treatment, suggesting that only the cytoophidium undergoing aggregation would be apparently perturbed by the drug. In order to confirm that the perturbation of cytoophidium assembly is due to the decrease of intracellular crowding, 25 mM, 50 mM and 100 mM sucrose was applied to the ACTD-treated cells to restore the crowding and cytoophidium assembly. While IMPDH failed to form detectable cytoophidia in most of ACTD treated cells in the medium with sucrose under 50 mM, nearly all cells were observed with cytoophidia in medium with 100 mM sucrose. (Figure 6D and E).

### The cytoophidium prolongs IMPDH half-life

Proteins located in intracellular inclusion bodies and other large aggregates, such as aggresomes and amyloids, are known to be more resistant to proteasomal degradation. In addition, CTPS cytoophidium formation can reduce CTPS ubiquitination and degradation by the proteasome in mammalian cells (Lin et al., 2018; Sun and Liu, 2019a). We suspected that the protection of component proteins from degradation is a common feature of different types of cytoophidia. To test this hypothesis, we aimed to compare the half-life of IMPDH protein in the cells with and without IMPDH cytoophidia.

HeLa cells were transfected with constructs encoding Myc-IMPDH2^WT^ and Myc-IMPDH^Y12A^, of which the expression is driven by a TRE-promoter. We harnessed the Tet-off system to manipulate the expression of the exogenous IMPDH2. Notably, the formation of cytoophidia could not be induced solely by IMPDH2 overexpression. In order to compare the half-life of exogenous IMPDH2 proteins in the cells with and without IMPDH cytoophidia, transfected cells were treated with MPA or DAU together with the doxycycline, which turns off the expression of exogenous IMPDH2. While MPA inhibits GTP biosynthesis and induces IMPDH cytoophidia, DAU inhibits CTP biosynthesis but does not induce IMPDH cytoophidia (Figure 7A). Thus, nucleotide synthesis and cell proliferation were impaired in both conditions but IMPDH cytoophidia should be present only in the culture with MPA. Western blotting was used to detect the remaining Myc-IMPDH2 in the treated cells (Figure 7B). While the level of Myc-IMPDH2 decreased by about 20% in MPA-treated Myc-IMPDH^WT^ expressing cells, levels of Myc-IMPDH2 dropped by about 60% in the other groups (Figure 7C). These results suggest that the half-life of IMPDH proteins is longer in cells with the cytoophidia, supporting the notion that the cytoophidium can protect component proteins from degradation.

**Figure 7.**
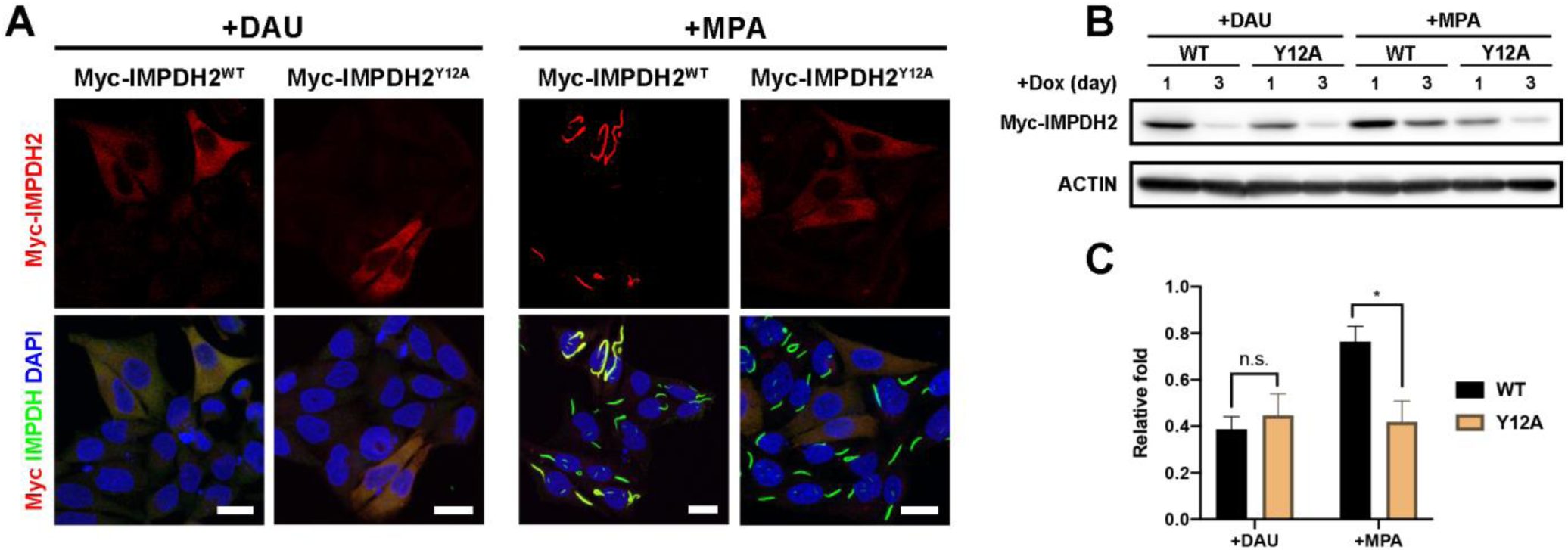
Formation of the cytoophidium prolongs the half-life of IMPDH2 protein. (A) Immunofluorescence of myc-IMPDH2^WT^ and myc-IMPDH2^Y12A^ overexpressing HeLa cells treated with DAU or MPA for 1 day. (B) Western blotting for the exogenous myc-IMPDH2 levels in transected HeLa cells treated with DAU or MPA for 1 and 3 days. (C) Quantitative data of the relative myc-IMPDH2 levels in the western blotting shown in (B). Scale bars = 20 μm in all panels. Error bars = S.E.M. Student’s *t* test, \**p* < 0.05.

## Discussion

The mechanisms and functions of protein aggregations have been intensively studied for decades. Many of them are known to comprise of unfolded or misfolded proteins and being associated with disorders and aging. For example, amyloid is a highly ordered fibrillar protein aggregate assembled by tau, prions and other proteins that feature the unique quaternary structure of β-sheets (Chiti and Dobson, 2017). Another well-known example, aggresomes, are dense inclusion bodies that sequester unfolded and misfolded proteins through highly regulated process (Kopito, 2000). In contrast, the cytoophidium is assembled by the aggregation of functional, correctly folded proteins and is normally reversible. Apart from being a storage spot of particular proteins, the cytoophidium is more likely an apparatus for regulating metabolic flux and other cellular functions. Therefore, the cytoophidium might be regarded as specialized protein aggregation, which serves as to regulate various functions of multivalent filament-forming proteins.

It has been demonstrated that the ER can regulate the dynamics of membraneless organelles like P-bodies and stress granules (Lee et al., 2020). At the early phase of cytoophidium assembly in cells, we find that IMPDH polymers form amorphous clumps, which would usually transform into compact filaments within minutes (Figure 3A and G). These IMPDH clumps are also associated with the ER and undergo fusion and fission (Figure 3C-F), suggesting that the ER may play a role in the formation and regulation of IMPDH cytoophidium.

Electrostatic interactions often require high protein concentrations and changes in salt or pH. The formation of multiple filamentous macrostructures in budding yeast have been shown to be induced by the change of pH, implying that the cytoophidium is assembled through the self-association of filamentous polymers (Hansen et al., 2021; Petrovska et al., 2014). Our findings support this notion as the presence of molecular crowders is sufficient to trigger purified human IMPDH2 proteins to reconstitute cytoophidium-like macrostructures (Figure 1). The negative charge at the loop^214-217^ of human IMPDH2 is possibly responsible for the interactions between polymers (Figure 2).

The effective concentration of interacting components is known to be critical to the establishment of interactions within protein aggregates. In some cases, the effective concentration of proteins could be modulated by expression levels and the crowding conditions of the cell. Yet, overexpression of CTPS and IMPDH is not sufficient to induce the cytoophidium in mammalian cells (Chang et al., 2018). In contrast, treatments of inhibitors such as DON and MPA could effectively induce CTPS and IMPDH cytoophidium assembly without changing expression levels of CTPS and IMPDH significantly (Keppeke et al., 2018; Lin et al., 2018). Since the polymers, but not oligomers, are the actual protomers of the cytoophidium, the concentration of protein polymers could be more important than their expression levels for cytoophidium assembly. Thus, the regulation of cytoophidium should be a combination of protein polymerization and the interaction between polymers, which could be regulated by ligand binding and the molecular crowding, respectively.

Approximately 40% of the cell volume is occupied by macromolecules, making the cytoplasm a crowded substance. Our data indicate that intracellular crowding plays an important role in the formation of the cytoophidium. In all tested cell lines, the numbers of detectable IMPDH cytoophidia were increased by the supplementation of sucrose in a dose-dependent manner (Figure 4). Conversely, the reduced molecular crowding by the inhibition of the synthesis of RNAs impaired the assembly of MPA-induced IMPDH cytoophidium (Figure 6). Ribosomal crowding has been demonstrated to be tuned by mTORC1 signaling pathway and can regulate the dynamics of various membraneless compartments (Delarue et al., 2018). These may partially explain the fact that the formation of IMPDH and CTPS cytoophidium is positively correlated with mTORC activity in eukaryotes (Andreadis et al., 2019; Chang et al., 2015; Sun and Liu, 2019b). Considering that the IMPDH and CTPS cytoophidium may function as an activity booster, this machinery may couple the metabolic and cellular status through the adjustment of molecular crowding.

When cells were treated with 300 mM sucrose or sorbitol, IMPDH, CTPS and PRPS simultaneously assembled into the cytoophidium (Figure 5D). These hyperosmotic conditions might be applied for a quick validation for the cytoophidium-forming capability of particular proteins. Although the regulation of the polymerization differs among proteins, such a universal condition for the simulation of cytoophidium assembly can provide a handy strategy for future studies.

On the other hand, the component proteins of dense protein aggregates such as amyloid and aggresome are known to be more resistant to the proteasomal degradation. Aggresomes are thought to be degraded through autophagy, one of the pathways by which large cellular structures are degraded (Garcia-Mata et al., 2002; Kopito, 2000). The degradation of amyloid, however, may require specialized proteases (Ries and Sastre, 2016). Previously, the formation of the cytoophidium has been reported to protect the CTPS from proteasomal degradation (Sun and Liu, 2019a). Consistently, we show that the cytoophidium can also prolong the half-life of IMPDH, suggesting this may be a common feature of the cytoophidium (Figure 7).

The physical state of biomolecular aggregates may determine certain properties of the compartment. For example, free diffusion of molecules may present in the liquid state, whereas in the gel and solid states, the molecular diffusion is more restricted. IMPDH cytoophidium is seemly a gel-like structure as the component proteins exhibited no flow in a previous fluorescence recovery after photobleaching (FRAP) analysis (Chang et al., 2018). In addition, a dense structure of the cytoophidium has been revealed with EM (Juda et al., 2014). These features implicate that the diffusion of molecules might be restricted in IMPDH cytoophidium. However, other evidences indicate that the occurrence of IMPDH assembly reflects the upregulation of GTP biosynthetic pathway or the increase of GTP consumption, suggesting that the cytoophidium is a catalytic active structure (Calise et al., 2018; Chang et al., 2015; Duong-Ly et al., 2018; Keppeke et al., 2018; Plana-Bonamaiso et al., 2020). Therefore, we propose that the architecture of the cytoophidium could act like a percolated system that allows the diffusion of its ligands, which are small molecules, but not of macromolecules. In other words, the large bundle of protein polymers may separate the interior proteins from other macromolecules while their catalytic reaction is less affected (Figure 8).

**Figure 8.**
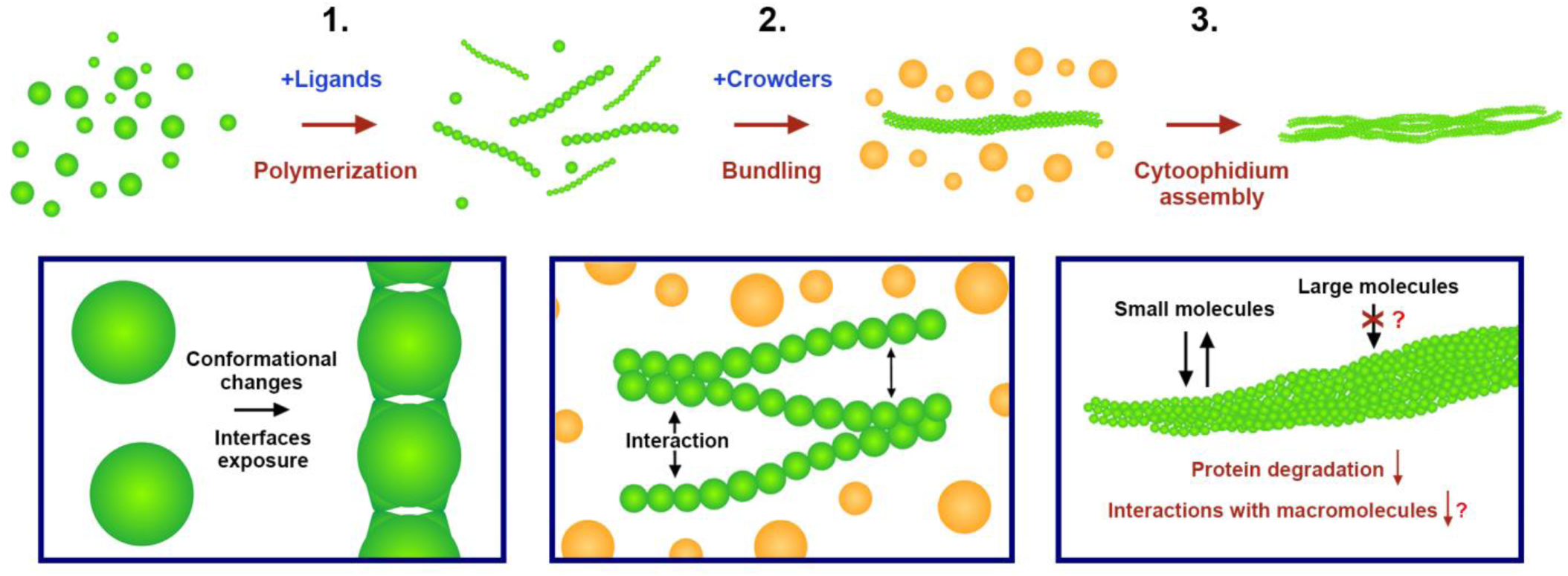
Illustration of the proposed molecular mechanism for the assembly of the cytoophidium. 1. Protein oligomers (octamer for IMPDH) polymerize into filamentous polymers under the regulation of conformational changes driven by the binding of ligands. Such conformational changes may expose or hide the interfaces responsible for the interactions between protomers. 2. Polymers self-associate into filament bundles through electrostatic interactions, which could be regulated by salt, pH, post-translational modifications, protein concentrations and molecular crowding. Such compact structures of filament bundles may allow small molecules percolate through the aggregate but restrict the interaction between macromolecules and component proteins. 3. Filament bundles may further assemble into larger structures, such as the cytoophidium.

In consistency with this model, the formation of CTPS cytoophidium was found to significantly attenuate the ubiquitination of CTPS (Sun and Liu, 2019a). In addition, the bundling of the translation initiation factor eIF2B filaments can downregulate general protein translation (Nuske et al., 2020). These could be led by reduced interactions between cytoophidium components and RNAs or proteins. IMPDH can act as a transcriptional and translational regulator in some models (Kozhevnikova et al., 2012; McLean et al., 2004; Mortimer et al., 2008). In human cancer cells, IMPDH has been shown to directly interact with other proteins including Y-box binding protein 1 (YB-1), Ras-related C3 botulinum toxin substrate 1 (RAC1), and circular RNAs (Bianchi-Smiraglia et al., 2021; Ruan et al., 2020; Wang et al., 2021). The regulation of these moonlight functions of IMPDH may be also correlated with the cytoophidium.

Taken together, in this study we clearly demonstrate that the bundling of IMPDH polymers is facilitated by molecular crowding and requires no other protein components. In mammalian cells, the assembly of IMPDH cytoophidium could be triggered by the increase of the concentration of IMPDH polymers and the intracellular crowding. The same mechanism might be projected to other cytoophidium-forming proteins. In addition, we show that the cytoophidium can prolong the half-life of IMPDH, suggesting a common function of different types of cytoophidia. Future researches are required to further explore each cytoophidium-forming protein and construct the general features of this growing protein family.

## Materials and Methods

### Cell culture and transfection

Human HEK 293T, HeLa, MCF7 and HCT116 cells were cultured in Dulbecco’s modified Eagle’s medium with high glucose, glutamine (SH30022.01, Hyclone) and supplemented with 10% FBS (Biological Industries) and 1% Penicillin-Streptomycin (60162ES76, YEASEN). Cells were kept in a 37 °C humid incubator with 5% CO2. The hyperosmotic medium was prepared with the supplementation of sucrose or sorbitol in the cultured medium with concentrations denoted in each experiment. MPA (M3536, Sigma-Aldrich), CX-5461 (HY-13323, Medchemexpress) and actinomycin D (SBR00013, Sigma-Aldrich), MPA (M3536, SigmaAldrich) and DAU (sc-394445, Santa Cruz Biotechnology) were supplemented into the culture medium in different experiments as described. Constructs encoding CMV-promoter driven OFP-hIMPDH2, GFP-IMPDH2, GFP-Sec61β, OFP-P2A-IMPDH2, OFP-P2A-IMPDH2^Y12A^, Flag-CTPS1, Flag-CTPS1^H355^ and TRE-promoter driven myc-IMPDH2 and myc-IMPDH2^Y12A^ were delivered into HEK 293T or HeLa cells with TurboFect™ transfection reagent (R0532, ThermoFisher). The expression of myc-IMPDH2 was turn off with culture medium containing 1 μg/ml doxycycline (ab141091, Abcam).

### Immunofluorescence

Cells were fixed with 4% paraformaldehyde in PBS for 10 minutes at room temperature. All antibodies were diluted in 1:500 dilution in PBS containing 2.5% BSA (A116563, Aladdin) and 0.25% Triton-X100 (X100, Sigma-Aldrich). Fixed samples were washed with PBS and incubated with primary antibody at room temperature for more than two hours. After washing with PBS, samples were then stained with secondary antibody at room temperature for about 2 hours. Mouse monoclonal anti-Flag (F1804, Sigma-Aldrich), mouse monoclonal anti-Myc (sc-40, Santa Cruz Biotechnology), rabbit polyclonal anti-IMPDH2 (12948-1-AP, ProteinTech), rabbit polyclonal anti-CTPS1 (15914-1-AP, ProteinTech), rabbit polyclonal anti-PRPS1 (15549-1-AP, ProteinTech), rabbit polyclonal anti-PAICS (GTX118341, GeneTex), rabbit polyclonal anti-GMPS (GTX114225, GeneTex), rabbit polyclonal anti-HPRT (GTX113466, GeneTex) Alexa Fluor 647-conjudated goat anti-mouse IgG (A21235, Invitrogen) and Alexa Fluor 488-conjugated donkey anti-rabbit IgG (A-21206, Invitrogen) antibodies were used.

### IMPDH cytoophidium in vitro reconstitution and negative staining

Same amount (1 μM) of purchased Human IMPDH2 recombinant protein (8349-DH-050, R&D systems) was incubated in 50 mM HEPES buffer (pH 8.5) containing 100 μM ATP and PEG-4000 at concentrations of 0, 100 or 200 mg/ml. The mixture was incubated at room temperature on with vertical shaking (200 r.p.m.) for 1 hour. The polymerization and bundling of IMPDH were analyzed with negative staining. Human IMPDH2 recombinant protein samples from the in vitro bundling procedures were loaded onto hydrophilic carbon-coated grids (400mech, Zhongjingkeyi Technology Co). The grids were then washed once with 0.1% uranium formate and dyed with 0.5% uranium formate. Imaging was performed on a 120 kV microscope (Talos L120C, ThermoFisher) with an Eagle 4 K x 4 K CCD camera system (Ceta CMOS, ThermoFisher).

### Live-cell imaging

OFP-IMPDH2 expressing HeLa cells on glass bottom chamber slides (C8-1.5H-N, Cellvis) with medium containing 100 μM MPA (M3536, Sigma-Aldrich), and maintained at room temperature when live-cell imaging was performed with a Leica SP8 STED 3X confocal microscope (Leica).

### 5-ethynyl uridine (EU) labelling

For EU incorporation, 30 min before fixation cells were incubated with 100 μM of EU (69075-42-9, Bidepharm). After 4% PFA fixation, a Click-iT™ (C10643, ThermoFisher) reaction was performed to bind Alexa Fluor 647 molecule to the EU incorporated to newly synthesized RNA. All procedures were performed according to the manufacturer protocol.

### Western blotting

Cell lysates were prepared with RIPA lysis buffer (20-188, Millipore) and quantitated for the amount of protein using a Bio-Rad Protein Assay Kit (5000002, Bio-Rad). Samples were run on a 12% polyacrylamide gel. PVDF membranes (GE Healthcare) were used for protein transfer. For immunolabelling, primary and secondary antibodies were incubated overnight diluted in PBST + 5% milk. Antibody labelling was revealed with SuperSignal West Pico Chemiluminescent Substrate (34579, ThermoFisher) and visualized in the chemiluminescence imaging system (GeneGnome XRQ, Syngene). Antibodies used: mouse anti-Myc monoclonal (9E10) antibody (1:1000, sc-40, SantaCruz); HRP-conjugated mouse monoclonal anti-ACTB antibody (1:3000, HRP-60008, ProteinTech). HRP-conjugated goal polyclonal anti-mouse IgG antibody (1:6000, 31430, ThermoFisher).

### Image analysis

All image-based quantification, including the number of nuclei and cytoophidia and fluorescence intensity, was analyzed with the software Fiji. The number of cell nuclei and percentage of cells with cytoophidia were counted manually. The quantification of the number of cytoophidia shown in figure 6C were performed with “analyze particles” and only particle size > 3 pixel^2^ were counted.

### Statistical analysis

Statistical analysis was performed in the software GraphPad Prism by using unpaired two-tailed Student’s *t*-tests or one-way ANOVA and Tukey’s test as denoted in the captions. All the quantification was from at least three repeats, and more than 100 cells were counted for each quantification. All error bars shown in graphs represent S. E. M.

## Supplementary information

**Figure S1.**
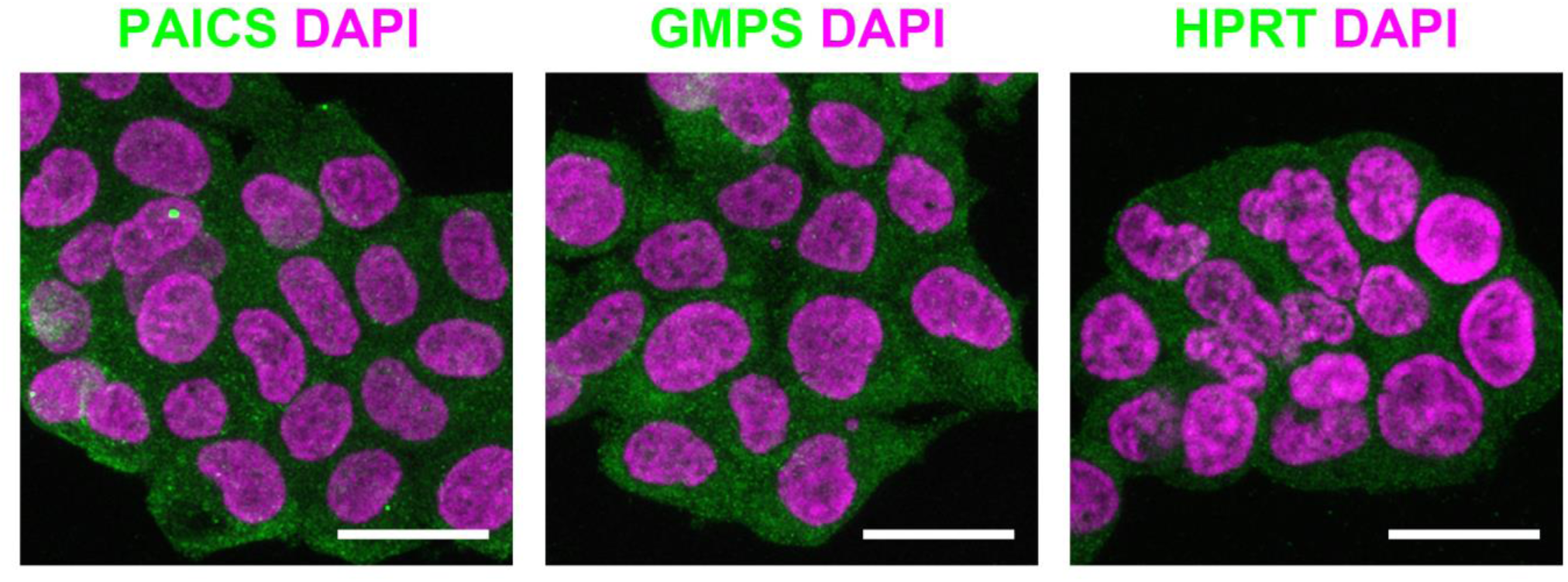
Hyperosmotic medium does not induce the aggregation of PAICS, GMPS and HPRT. Immunofluorescence of wildtype HEK 293T cells treated with 300 mM sucrose for 3 hours. No detectable aggregates of PAICS, GMPS or HPRT were observed in the cells. Scale bars = 20 μm.

**Figure S2.**
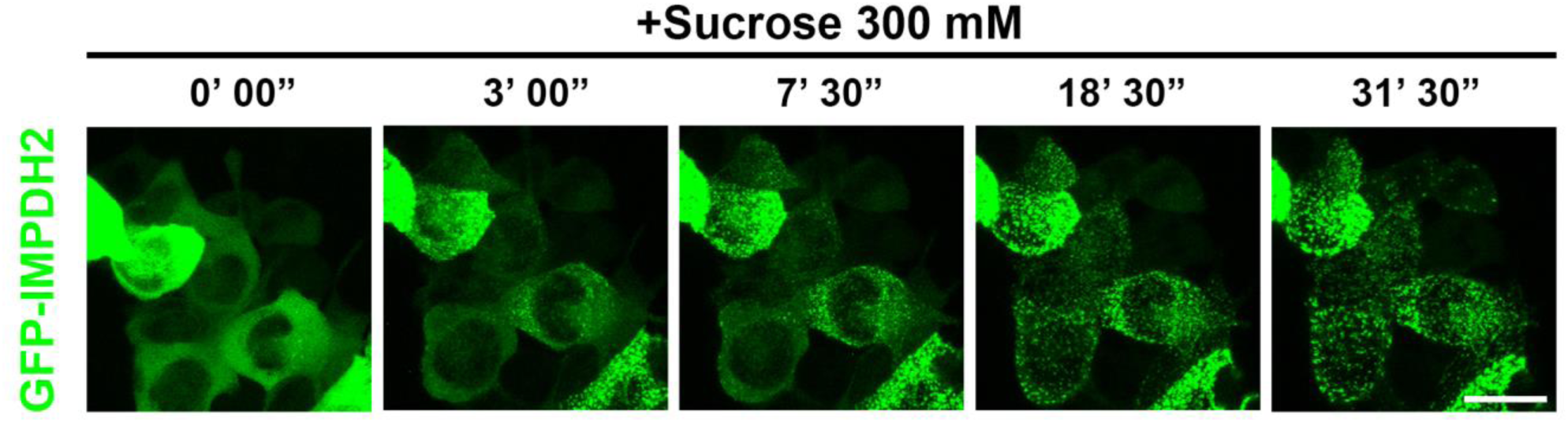
The aggregation of GFP-IMPDH2 under the treatment of hyperosmotic medium. Representative frames of live-cell imaging of GFP-IMPDH2 expressing HEK 293T cells treated with sucrose (300 mM). Scale bars = 20 μm.

**Movie S1. Live-cell imaging showing the dynamics of IMPDH amorphous clumps.** OFP-IMPDH2 expressing HeLa cells were treated with MPA. Time intervals of each frame is 40 seconds.

**Movie S2. Live-cell imaging showing an IMPDH amorphous clump transforms in association with the ER.** GFP-Sec61β (green) and OFP-IMPDH2 (red) expressing HeLa cells were treated with MPA. Time intervals of each frame is 5 seconds.

**Movie S3. Live-cell imaging showing the transformation of IMPDH amorphous clumps to filaments.** OFP-IMPDH2 expressing HeLa cells were treated with MPA. Time intervals of each frame is 60 seconds.

